# Spatio-relational inductive biases in spatial cell-type deconvolution

**DOI:** 10.1101/2023.05.19.541474

**Authors:** Ramon Viñas, Paul Scherer, Nikola Simidjievski, Mateja Jamnik, Pietro Liò

**Affiliations:** Department of Computer Science and Technology, University of Cambridge, Cambridge, UK

## Abstract

Spatial transcriptomic technologies profile gene expression *in-situ*, facilitating the spatial characterisation of molecular phenomena within tissues, yet often at multi-cellular resolution. Computational approaches have been developed to infer fine-grained cell-type compositions across locations, but they frequently treat neighbouring spots independently of each other. Here we present GNN-C2L, a flexible deconvolution approach that leverages proximal inductive biases to propagate information along adjacent spots. In performance comparison on simulated and semi-simulated datasets, GNN-C2L achieves increased deconvolution performance over spatial-agnostic variants. We believe that accounting for spatial inductive biases can yield improved characterisation of cell-type heterogeneity in tissues.

## 1. Introduction

Analysing the spatial organisation of cells within a tissue can shed light on fundamental biological processes, including intercellular communication (Fischer et al., 2023) and organogenesis (Lohoff et al., 2022), and mechanisms of diseases like cancer, diabetes, and autoimmune disorders (Solinas et al., 2007; Wang et al., 2013; Vlahopoulos et al., 2015). Spatial transcriptomics technologies have recently enabled gene expression profiling *in situ*, but they often lack single-cell resolution, impeding fine-grained characterisation of cellular heterogeneity and effective reconstruction of tissue architectures.

Computational approaches for cell-type deconvolution in spatial transcriptomics offer a scalable solution to these challenges. These strategies often identify resident cell types from the RNA sequencing of dissociated single cells, yielding cell-type-specific gene expression signatures, and then infer the cell-type composition of every profiled spot (Elosua-Bayes et al., 2021; Cable et al., 2022; Dong & Yuan, 2021). A cutting-edge method in this family is Cell2Location (Kleshchevnikov et al., 2022), a Bayesian deconvolution approach that captures cell-type relationships through a hierarchical model and handles technical sources of variation like differences in mRNA detection sensitivity. Despite numerous benefits, however, existing deconvolution approaches treat spots independently of each other.

In this study, we investigate whether incorporating spatio-relational information leads to improved cell-type mapping. Building on the observation that neighbouring spots often exhibit similar cell-type compositions (Figure 1), we extend Cell2Location (C2L) to incorporate spatial inductive biases. Our approach, named GNN-C2L, propagates learnable messages on the proximity graph of spot transcripts, effectively leveraging the spatial relationships between spots and exploiting the co-location of cell-types (Figure 2). We conduct an extensive ablation study on synthetic and real spatial transcriptomics datasets and show improved deconvolution performance of GNN-C2L over spatial-agnostic variants. Altogether, our work leverages proximal inductive biases to facilitate enhanced reconstruction of tissue architectures.

**Figure 1:**
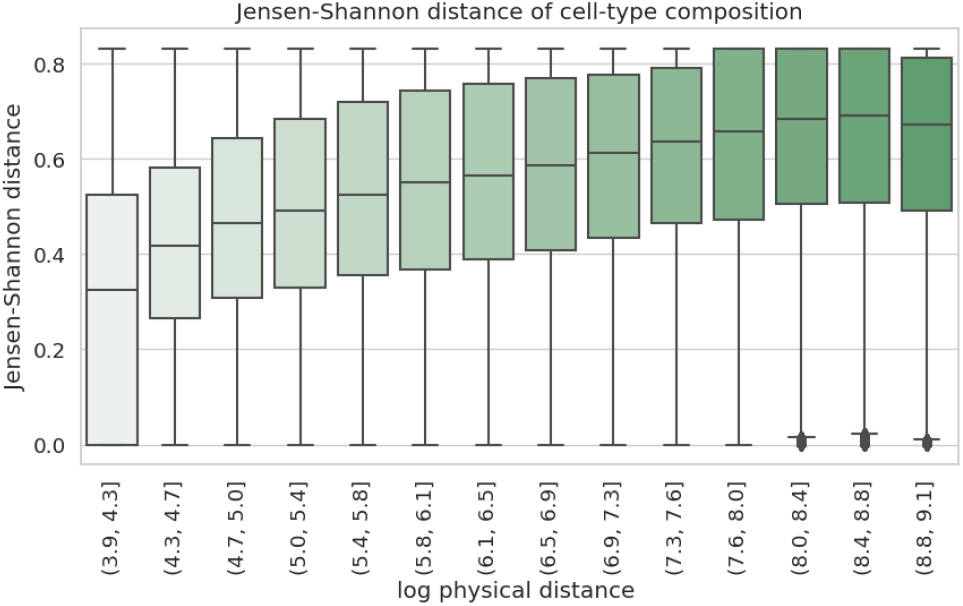
Jensen-Shannon distance of cell-type proportions by spot distance in Xenium dataset (breast cancer, convolved spots of size 50μm). Closer spots tend to exhibit similar cell-type composition.

**Figure 2:**
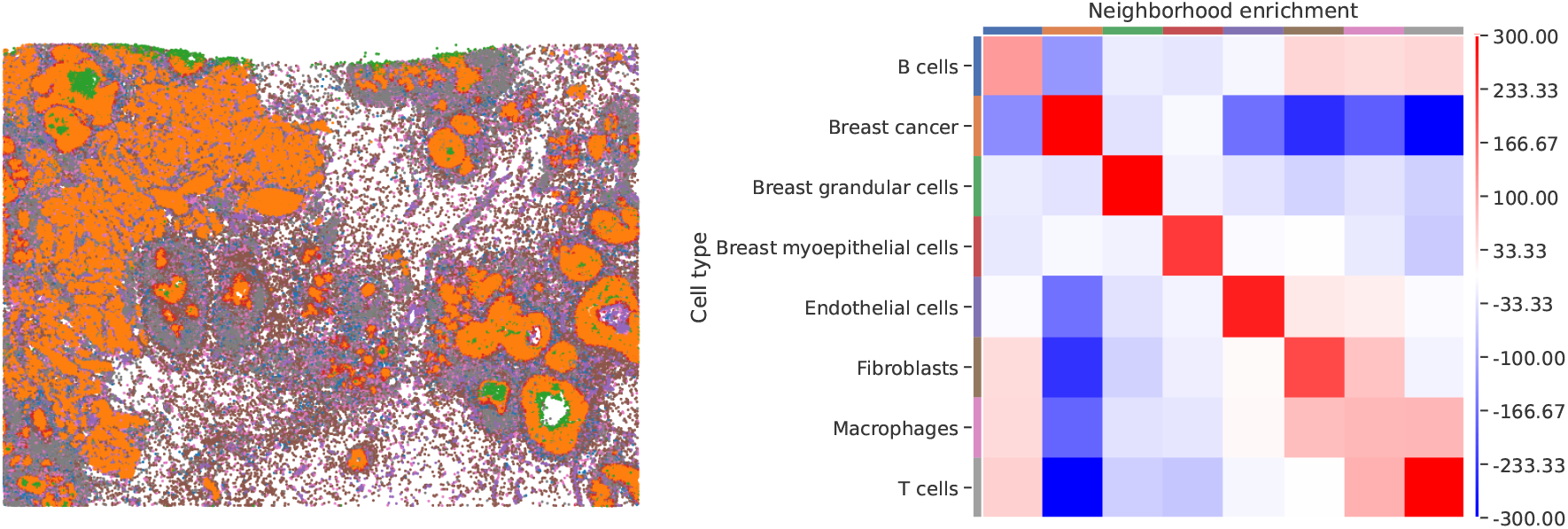
Neighbourhood enrichment analysis on the Xenium dataset (breast cancer). The color legend is given by the y-axis of the neighbourhood heatmap. (*left*) Spatial transcriptomics data colored by cell-type. (*right*) Neighbourhood enrichment z-scores (red and blue indicate enrichment and depletion in the neighbourhood of nearest neighbours, respectively). Cells from the same cell-type tend to co-locate (e.g. breast cancer cells). Immune cells — including T cells, B cells, and macrophages — work in conjunction to modulate the anti-cancer immune response (Gonzalez et al., 2018). Utilising relational inductive biases could therefore enhance the effectiveness of spatial deconvolution models, thereby improving the characterisation of tumor microenvironments at different stages of cancer progression.

## 2. Methodology

### Problem formulation

Let ***D*** ∈ ℝ^*S*×*G*^ denote a count matrix of RNA reads captured at *S* spots for *G* genes, using one or multiple batches (e.g. 10x Visium slides or Slide-seq pucks). Let *d*_*s,g*_ be the entry of this matrix with the number of reads for gene *g* in spot *s*. Let ***C*** ∈ℝ^*F* ×*G*^ denote a matrix of *F* reference cell-type signatures for the same set of *G* genes (e.g. these signatures can be obtained from dissociated single-cell RNA-seq, Appendix A). Denote by *c*_*f,g*_ the expression of gene *g* in signature *f*. Given the count matrix ***D*** and cell-type signatures ***C***, our goal is to infer the cell-type composition ***X*** ∈ ℝ^*S*×*F*^ of every spot.

### Cell2Location

Our relational approach builds on Cell2Location (Kleshchevnikov et al., 2022, Appendix A), which models the per-spot read counts ***D*** as Negative Binomial (NB) distributed:

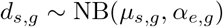

where *α*_*e,g*_ is an experiment-and gene-specific over-dispersion parameter and the unobserved expression rate *μ*_*s,g*_ is modelled as a linear function of the reference cell-type signatures *c*_*f,g*_:

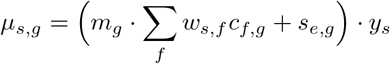

where *w*_*s,f*_ corresponds to the abundance of cell-type *f* at location *s, m*_*g*_ is a scaling parameter specific to gene *g, s*_*e,g*_ is an experiment-and gene-specific additive shift, and *y*_*s*_ is the detection sensitivity at spot *s*.

### GNN-C2L

We propose a hierarchical model for cell-type composition inference that incorporates proximal relationships between spots. Let *𝒩*(*s*) be the set of neighbor indices for spot *s*. This set of neighbours can be adapted to various spatial arrangements (e.g. hexagonal neighbourhoods for 10X Visium data) and k-hop neighbourhoods. To account for the neighbourhood information, we introduce a latent variable *γ*_*s,f*_ representing the neighbour-aware cell-type abundances:

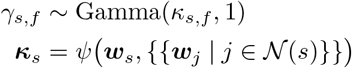

where the shape parameter *κ*_*s,f*_ depends on the latent variables ***w***_*s*_ and {{***w***_*j*_| *j* ∈ *𝒩* (*s*)}} of spot *s* and its neighbours through a transformation *ψ*(·). Unlike Cell2Location, this effectively adds graphical dependencies between the neighbour-informed variables *γ*_*s,f*_ and the latent variables *w*_*s,f*_ (Appendix A). Importantly, computing *γ*_*s,f*_ as a function of ***w***_*s*_ allows capturing cell-type co-location patterns.

We then compute mean parameter *μ*_*s,g*_ of the Negative Binomial NB(*μ*_*s,g*_, *α*_*e,g*_) likelihood using the neighbour-aware cell-type abundances *γ*_*s,f*_ :

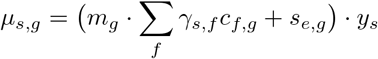

For all parameters, we utilise the validated hierarchical priors and hyperpriors of Cell2Location (Appendix A).

### Incorporating spatial inductive biases

The form of *ψ*(·) determines the inductive biases of the model. In this study, we construct a proximity graph of spatially localised spots, i.e. we consider physically adjacent spots, allowing for different spatial arrangements (e.g. hexagonal neighbourhoods for 10X Visium data) and k-hop neighbourhoods. We consider several graph neural network architectures for *ψ*(·), starting with simple graph convolutional network (Wu et al., 2019) to validate whether homophily (enacted by feature propagation) is a useful inductive bias, and introducing other GNN operators to allow for a more expressive use of the available spatio-relational data. We also consider a standard multi-layer perceptron as a baseline to assess whether performance changes can be attributed to similarly parametrised spatial-agnostic transformations. We next describe the alternatives for *ψ* in greater detail.

### MLP-C2L

As a spatial-agnostic control, we model *ψ*(·) with an MLP, i.e. ***κ***_*s*_ = MLP(***w***_*s*_), using a softplus activation function. This model does not utilise any spatial relationships between the spots and, alongside Cell2Location, serves as a control for our hypothesis.

### SGC-C2L

We construct a GNN-C2L variant using Simple Graph Convolutional (SGC) layers (Wu et al., 2019; Scherer et al., 2019). Let *d*_*s*_ =| *𝒩* (*s*) | be the node degree of spot *s*. A SGC layer computes the neighbour-aware features ***κ***_*s*_using a weighted average of the latent variables ***w***_*s*_ in the local neighbourhood:

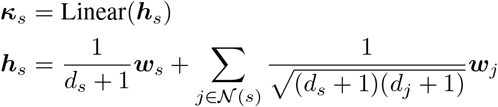

The feature propagation mechanism biases the representations ***κ***_*s*_ of neighbouring spots to become more similar to each other, using a degree-normalised adjacency matrix with self-loops. Thus, this simple MLP extension encourages homophilous latent cell-type distributions. Optionally, we can apply an activation function after the linear transformation and stack several SGC layers to expand the receptive field.

### GAT-C2L

We increase the expressivity of *ψ*(·) by utilising graph attention networks, specifically GATv2 (Brody et al., 2022). Unlike the constant, degree-dependant neighbouring contribution in the SGC-C2L model, the GATv2-C2L variant employs a learnable attention mechanism with increased control of contribution strengths, allowing to capture both homophilic and cell-type co-location patterns:

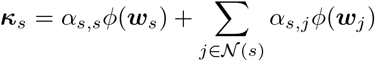

where *ϕ* is an MLP with a softplus activation function. We define the attention coefficient *α*_*s,j*_ as:

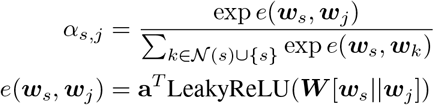

where ‖ is the concatenation operation and **a** and ***W*** are learnable parameters shared across spots, allowing the neural network to mix signals over the different cell types.

### Training and inference

We approximate the model parameters through variational inference. For every latent variable, we use a univariate normal distribution to approximate the posterior and utilise a softplus activation to ensure a positivity. Minimisation of the ELBO jointly trains the parameters of the model (and the incorporated GNNs) as well as the variational distribution. After optimisation, we estimate the cell-type abundances of every spot *s* by averaging *γ*_*s,f*_ over 1000 samples of the variational distribution.

## 3. Experimental setup

We study whether incorporating spatial relationships via graph neural networks leads to enhanced cell-type mapping.

### Datasets

To quantitatively benchmark the baselines, we utilised a synthetic dataset introduced in Cell2Location (Kleshchevnikov et al., 2022) for which we knew the “true” cell-type abundances of each spot. The construction of this dataset is detailed extensively in (Kleshchevnikov et al., 2022). Moreover, we evaluated the methods using two real datasets, MPOA (Moffitt et al., 2018) and Xenium (breast cancer) (Janesick et al., 2022), that have single-cell resolution (yet fewer genes are profiled). To simulate real spots, we divided the tissues into squared spots of size 100*μ*m and summed the expression of all cells within every square (Appendix B). For the Xenium and MPOA datasets, we constructed the cell-type signatures by averaging the read counts of all cells from every given cell-type (Appendix C).

### Hyperparameter settings

We used the same hyperparameters for every baseline where applicable. We set the hidden dimensions of each layer to 64 and used a single GNN layer (i.e. 1-hop receptive field). We conducted an ablation study using more graph layers in Appendix D. We minimised the variational lower bound using Adam (Kingma & Ba, 2014) with learning rate of 0.001 for 25,000 epochs in all datasets.

### Evaluation metrics

For all datasets we assessed performance using the average Pearson *R* correlation, Jensen-Shannon Divergence (JSD), and the area under precision-recall curve (AUPRC) (macro-averaged over cell-types) between the ground-truth and inferred cell-type proportions. We computed Pearson *R* over the flat ground-truth and inferred cell-type proportions. We calculated the Jensen-Shannon Divergence between the per-spot ground-truth and inferred cell-type proportions. We binarised the true cell abundance matrix to show which cell types were present in which locations, and then used the inferred cell-type proportions to compute the AUPRC.

## 4. Results and discussion

We benchmark spatial-agnostic (C2L, MLP-C2L) and spatial-aware (SGC-C2L, GAT-C2L) baselines on simulated and semi-simulated (MPOA and Xenium) spatial transcriptomics data (Table 1).

**Table 1:** Average Pearson *R*, Avg. Jensen Shannon divergence (JSD), and AUPRC scores and standard deviation of 5 seeded runs of each model over all spots. For the synthetic dataset, scores for subcategories of cell types exhibiting distinct cell abundance patterns are also provided. Bold numbers indicate best-performing method for each category of cell types being evaluated for each metric. Overall, the GNN-C2L spatial-aware variants attained equal or superior deconvolution scores than spatial-agnostic baselines.

| Method | Simulated |  |  |  |  | Semi-simulated |  | Metric |
| --- | --- | --- | --- | --- | --- | --- | --- | --- |
|  | ALL | UHCA | ULCA | RHCA | RLCA | MPOA | Xenium |  |
| C2L | 0.683 ± 0.002 | 0.882 ± 0.001 | 0.519 ± 0.007 | 0.836 ± 0.004 | 0.422 ± 0.003 | 0.929 ± 0.001 | 0.928 ± 0.002 | $R$ |
| MLP-C2L | 0.672 ± 0.024 | 0.866 ± 0.008 | 0.661 ± 0.021 | 0.865 ± 0.007 | 0.404 ± 0.040 | 0.920 ± 0.005 | **0.929 ± 0.000** | |
| SGC-C2L | 0.699 ± 0.023 | 0.876 ± 0.008 | **0.708 ± 0.020** | 0.883 ± 0.006 | 0.439 ± 0.041 | 0.936 ± 0.001 | 0.928 ± 0.000 | |
| GAT-C2L | **0.737 ± 0.013** | **0.885 ± 0.018** | 0.695 ± 0.032 | **0.888 ± 0.004** | **0.492 ± 0.032** | **0.936 ± 0.001** | 0.928 ± 0.000 | |
| C2L | 0.468 ± 0.001 | **0.202 ± 0.002** | 0.496 ± 0.001 | 0.421 ± 0.002 | 0.509 ± 0.001 | 0.204 ± 0.001 | 0.213 ± 0.005 | Avg. JSD |
| MLP-C2L | 0.457 ± 0.006 | 0.230 ± 0.012 | 0.473 ± 0.007 | 0.387 ± 0.006 | 0.503 ± 0.009 | 0.199 ± 0.004 | 0.211 ± 0.001 | |
| SGC-C2L | 0.446 ± 0.006 | 0.224 ± 0.011 | 0.460 ± 0.007 | **0.368 ± 0.005** | 0.493 ± 0.009 | 0.189 ± 0.001 | **0.211 ± 0.001** | |
| GAT-C2L | **0.435 ± 0.003** | 0.209 ± 0.021 | **0.458 ± 0.014** | 0.369 ± 0.001 | **0.482 ± 0.006** | **0.188 ± 0.001** | 0.212 ± 0.000 | |
| C2L | 0.591 ± 0.003 | 0.932 ± 0.006 | 0.477 ± 0.005 | 0.783 ± 0.003 | 0.591 ± 0.003 | **0.956 ± 0.001** | 0.873 ± 0.003 | AUPRC |
| MLP-C2L | 0.675 ± 0.002 | 0.963 ± 0.006 | 0.590 ± 0.004 | 0.804 ± 0.004 | 0.675 ± 0.002 | 0.951 ± 0.001 | 0.883 ± 0.001 | |
| SGC-C2L | 0.719 ± 0.002 | 0.977 ± 0.004 | 0.646 ± 0.006 | **0.861 ± 0.001** | 0.719 ± 0.002 | 0.955 ± 0.000 | 0.884 ± 0.000 | |
| GAT-C2L | **0.722 ± 0.002** | **0.978 ± 0.004** | **0.664 ± 0.004** | 0.858 ± 0.003 | **0.722 ± 0.002** | 0.952 ± 0.001 | **0.884 ± 0.000** | |

### Results on simulated dataset

We studied deconvolution performance on the synthetic data over: 1) **ALL**: all cell types, 2) ubiquitious high cell abundance (**UHCA**): 3 high-abundance cell types spatially distributed in uniform manner across the tissue, 3) ubiquitious low cell abundance (**ULCA**): 5 low-abundance cell types spatially distributed in uniform manner across the tissue, 4) regional high cell abundance (**RHCA**): 9 cell types with local distribution patterns, that is, cell types are clustered in specific locations with high abundance and exhibit 0 abundance elsewhere, 5) regional low cell abundance (**RLCA**): 32 low-abundance cell types that have local distribution patterns.

Overall, GNN-C2L consistently outperformed the spatial-agnostic baselines on the synthetic data (Table 1). We observed a marked increase in performance through the utilisation of proximal relations across different metrics and subtasks. Spatial-aware baselines achieved the best scores in 13 out of 15 cases, especially for cell types with low cell abundance (ULCA and RLCA). The performance difference was particularly apparent from the overall scores of MLP-C2L (ALL R: 0.672±0.024, JSD: 0.457±0.006, AUPRC: 0.675±0.002) and SGC-C2L (ALL R: 0.699±0.023, JSD: 0.446±0.006, AUPRC: 0.719±0.002) — both baselines utilised the number amount of learnable parameters, yet only SGC-C2L propagates information across spots. It is balso worth noting that using additional parameters may result in degraded performance, i.e. compared to C2L (ALL *R* : 0.683±0.002), MLP-C2L attained reduced Pearson *R* correlation and increased variance (ALL *R* : 0.672±0.024). Altogether, our results highlight the superior ability of GNN-C2L to perform cell-type deconvolution.

### Results on semi-simulated datasets

In performance comparison on the semi-simulated datasets (MPOA and Xenium), the spatial-aware GNN-C2L variants achieved equal or better deconvolution performance than the spatial-agnostic baselines (Table 1). On MPOA, all baselines performed well — it is worth noting that this is a considerably smaller dataset with larger spot sizes (per-spot average of 18 cells) compared to the synthetic (∼9 cells per spot) and Xenium (∼10 cells per spot) datasets. This may have an effect on the specificity of the transcript readings as well as the usefulness of local information considering the size of micro-architectures in the tissue. We observed that GAT-C2L had the best scores in 2 out of 3 metrics (R: 0.492±0.032, JSD: 0.188±0.001), while Cell2Location was superior in terms of AUPRC (0.956±0.001). In the Xenium dataset, all baselines attained comparable results (e.g. C2L R: 0.928 ± 0.000, GAT-C2L R: 0.928 ± 0.000; C2L AUPRC: 0.873 ± 0.003, MLP-C2L AUPRC: 0.883 ± 0.001, SGC-C2L AUPRC: 0.884 ± 0.000).

Collectively, our results suggest that spatial deconvolution can benefit from spatio-relational inductive biases, with potential for enhanced reconstruction of tissue architectures.

## Code availability

GNN-C2L is publicly available at https://github.com/paulmorio/GNN-C2L.

## A. Cell2Location and GNN-C2L model description

In order to describe our model we will first go over the construction of Cell2Location (Kleshchevnikov et al., 2022). Cell2Location is a Bayesian inference model built in a hierarchical manner to account for different sources of confounding experimental information.

Let *D*∈ ℝ^*S*×*G*^ denote a mRNA count matrix with its entries corresponding to mRNA count at spot *s*∈ {1, …, *S*} from one or multiple batches (i.e. 10x Visium slides or SlideSeq pucks) for genes *g* ∈ {1, …, *G*}. Let *𝒞* ∈ ℝ^*F* × *G*^ denote a matrix of reference cell type signatures obtained from learning on the scRNA data (see Section A.1.1). Note that *𝒟* and *𝒞* need to be aligned such that they cover the same set of genes *G*. Cell2Location models the elements of *D* as Negative Binomial distributed (NB), given an unobserved expression level (rate) *μ*_*s,g*_ and a geneand experimentspecific over-dispersion parameter *α*_*e,g*_:

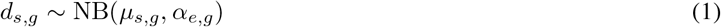

This can be equivalently expressed as a Gamma-Poison mixture with a Poisson likelihood (count measurement model) and a Gamma-distributed mean (expression dispersion model):

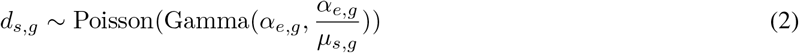

The expression level of genes *μ*_*s,g*_ in the mRNA count space is modelled as a linear function of the reference cell type expression signatures:

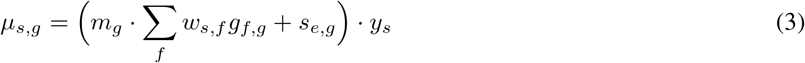

- Here *w*_*s,f*_ denotes regression weight of each reference signature *f* at location *s*, which can be regarded as the abundance or proportion of cells expressing reference cell type signature *f* at *s*. This is the latent variable that we care about and intend to infer.
- *m*_*g*_ denotes a gene-specific scaling parameter, which adjusts for global differences in expression estimates between technologies.
- *s*_*e,g*_ captures gene specific additive shift (due to free-floating RNA).
- *y*_*s*_ denotes a location-specific scaling parameter, which models variation in RNA detection sensitivity across locations and experiments. This parameter scales the contributions of the cell types and the gene specific additive shift *s*_*e,g*_.

We dive into the derivation of the prior distributions for each of the latent variables.

### Cell abundance across locations

*w*_*s,f*_ This is Gamma distributed according to

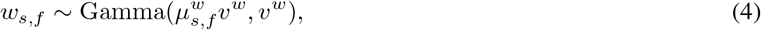

where *v*^*w*^ is a fixed hyperparameter denoting prior strength, the prior mean parameter is modelled in a hierarchical fashion, decomposing the regression weights into *R* latent groups of cell types *r* = {1, …, *R*} (by default Cell2Location uses *R* = 50) accounting for linear dependencies in spatial abundance of cell types:

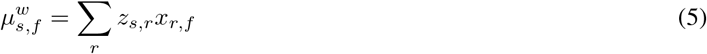

Intuitively, *R* can be considered as the number of cellular compartments or zones in the tissue that are characterised by shared cell type composition. The authors observed that the sensitivity of mapping cell types with small transcriptional difference increases when accounting for these dependencies. For our purposes we are not interested in this parameter and utilise the default for all experiments.

*z*_*s,r*_ and *x*_*r,f*_ are prior distributions defined to control absolute scale of the cell type abundance estimates, guiding *w*_*s,f*_ to the scale of the number of the abundance of cells expressing reference cell type signature *f* at location *s*. These priors are important because there is a non-identifiability between *m*_*g*_, *y*_*s*_, *w*_*s,f*_ unless informative priors are constructed for each of them. Moreover, the prior distributions help control the sparsity of how many cell types *f* are expected at each spot *s*, facilitating application of Cell2Location to tissues and technologies with varying numbers of cells and cell types per location. The hyperparameters controlling the *w*_*s,f*_ prior can be estimated from a paired histology image or a literature based estimate. The hyperparameters controlling *y*_*s*_ can be estimated based on total RNA counts in the input data and the quality of the experiment. The prior distribution *z*_*s,r*_ is defined as follows:

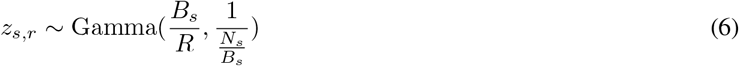

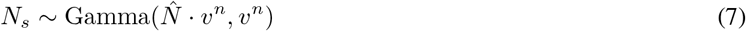

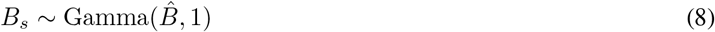

where *N*_*s*_ is associated to the latent average number of cells in each location, and *B*_*s*_ is the latent number of groups *r* expected in each spot *s*. 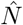 is a user-defined estimate of the expected number of cells per location (see end of this section). 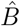 is the expected average number of cellular components or zones per location; by default it is initialised to 7. *v*^*n*^ denotes a prior strength. The construction is done such that ∑ _*r*_ *z*_*s,r*_≥ *N*_*s*_. In other words, the expectation of the sum over *z*_*s,r*_ equals the expected number of cells per location *N*_*s*_ and that on average each location has a high value of *z*_*s,r*_ for *B*_*s*_ expected cell type groups.

*x*_*r,f*_ represents the contribution of each latent cell type group *r* to the abundance of each cell type *f* and is Gamma distributed in the following manner:

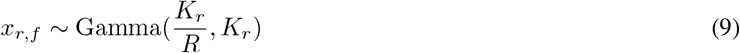

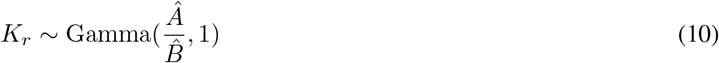

*K*_*r*_ represents the unobserved number of cell types for each group *r*. This prior controls the absolute values of *x*_*r,f*_ such that on average ∑ _*r*_ *x*_*r,f*_= 1. *Â* and 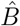 intuitively represent the expected number of cell types per spot, and the expected number of cellular components per spot respectively. By default both *Â* and 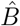 are initialised at 7, indicating the prior belief that the spatial abundance of each cell type *f* is independent from other cell types. Conversely, each group *r* has have a large value of *x*_*r,f*_ for many cell types *f* when 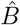.

### Gene specific multiplicative scaling factor *m*_*g*_

This is modelled as Gamma distributed with hierarchical prior *μ*^*m*^ and *α*^*m*^ which provide regularisation:

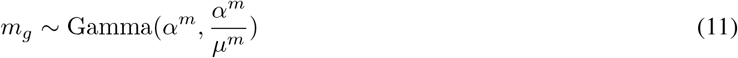

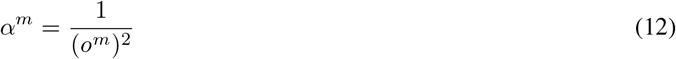

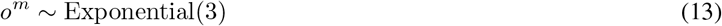

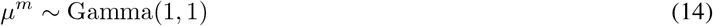

### The prior on detection efficiency *y*_*s*_ per location

This prior is selected to discourage over normalisation, such that unless data has evidence of strong within-experiment variability in RNA detection sensitivity across locations, it is assumed to be small and close to the mean sensitivity for each experiment or batch *y*_*e*_:

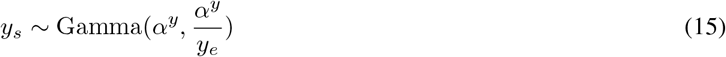

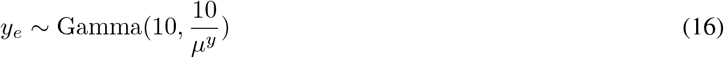

where *α*^*y*^ is a user defined hyperparameter that regularises within experiment variation; and *y*_*e*_ is a latent detection efficiency for each batch or experiment *e. mu*^*y*^ is estimated using observed variables (*d*_*s,g*_ and *g*_*f,g*_) and the hyperparameter 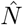 in the following manner:

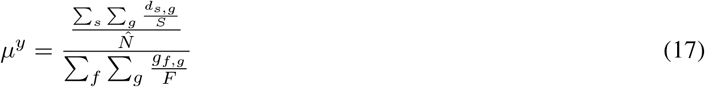

where we remind ourselves that *S* is the total number of spots, and *F* is the total number of cell types.

### Overdispersion containment prior

*α*_*e,g*_ A containment prior (Simpson et al., 2017) is used to model the latent variance of the negative binomial distribution modelling *d*_*s,g*_. This prior is intended to encourage simplicity of the NB model making it closer to the Poisson distribution (via larger *α*_*e,g*_ values producing bigger probability masses). Thus the prior expresses a belief that most genes have low overdispersion:

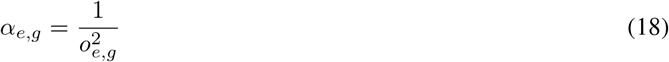

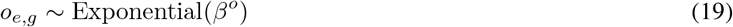

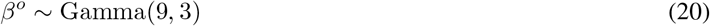

where the constants 9 and 3 correspond to values of *α*_*e,g*_ observed in previous modelling studies (Kleshchevnikov et al., 2022; Lopez et al., 2018).

### Additive shift bias *s*_*e,g*_

This latent variable accounts for confounding effects on RNA counts for every gene *g* for every experiment *e* caused by phenomena such as free-floating RNA in the tissue sample. This additive shift is modelled using a Gamma distribution again with hierarchical experiment specific priors 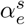 and 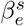which provide regularisation:

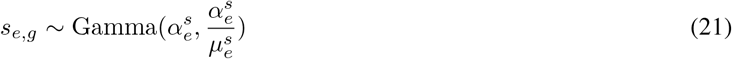

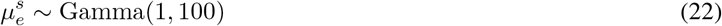

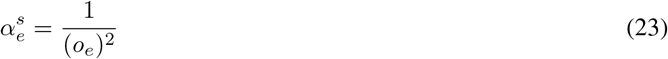

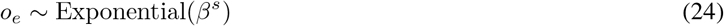

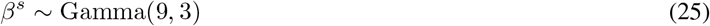

The hierarchical priors 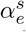 and 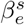 model the variation across experiments *e. β*^*s*^ serves as a hyperparameter that allows the model to learn 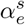 rather than requiring a user to define it.

This leaves Cell2Location with two hyperparameters whose values have to be considered based on the dataset and how the spatial transcriptomics experiment was performed:

1. 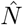: the expected number of cells per spot. This is a tissue-level global estimate, which can be derived from paired histology images (see the H&E stained image in Figure 3). An estimate may be obtained by manually counting nuclei in a set of random spots using the appropriate software from the measurement device (e.g. 10x Loupe Browser for the Visium slides outputs). When this is not available one can also use the size of the captured regions relative to an expected cell size.
2. *α*^*y*^: the regularising hyperparameter for within-experiment variation of RNA detection sensitivity. In the default setting it is assumed that there is little variability in the RNA detection sensitivity so *α*^*y*^ is set to *α*^*y*^ = 200 which results in values of *y*_*s*_ close to the mean sensitivity for each experiment *y*_*e*_. A lower value would enforce a stronger normalization to the sensitivity, and a correspondingly lower regularisation toward the mean sensitivity.

**Figure 3:**
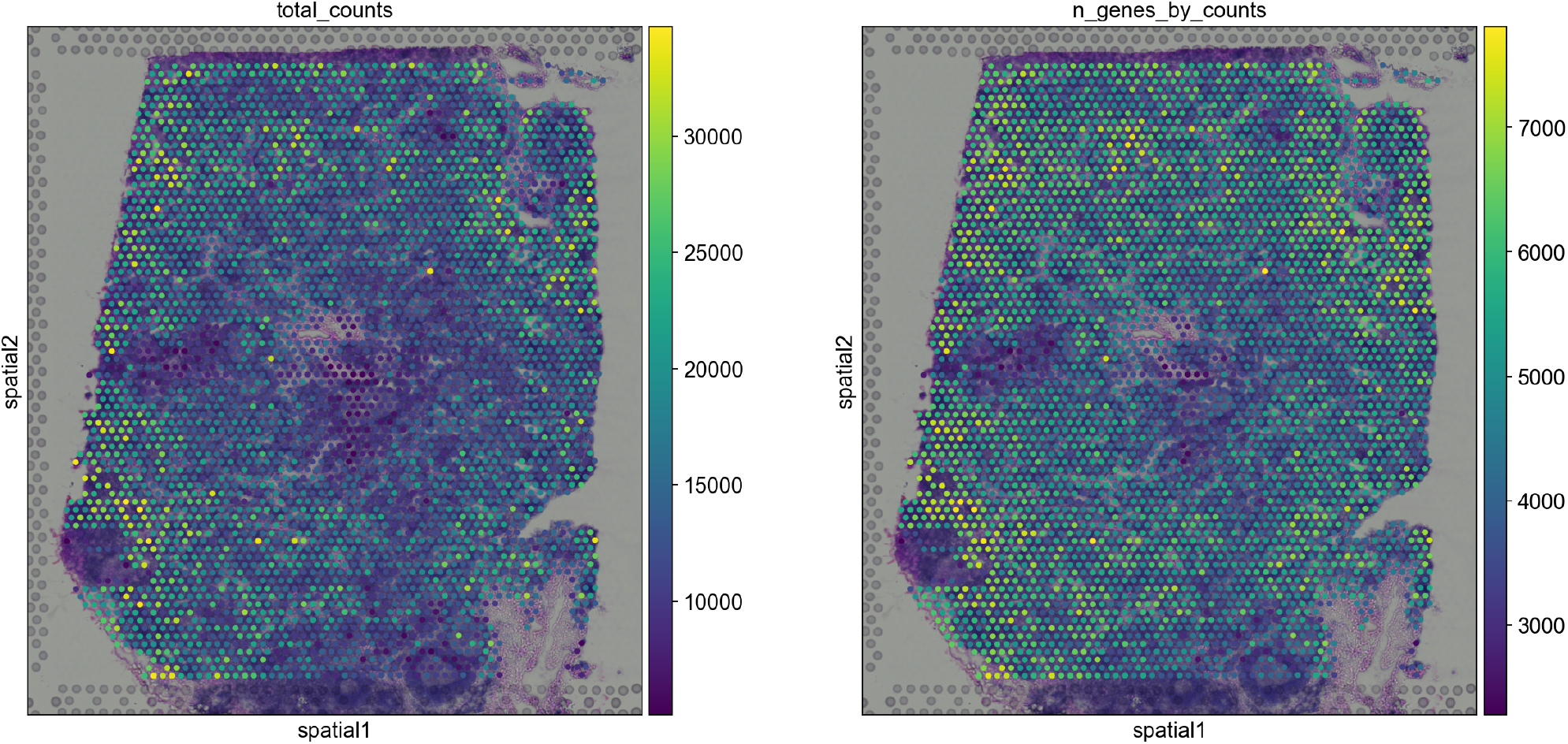
Visium slide of a human lymph node tissue publicly available at the 10x Genomics dataset portal, accessed through scanpy (Wolf et al., 2018). In these images the circular spots are overlaid over the Hematoxylin and eosin stain (H&E) image of the tissue sample. Note the non-uniform dispersion of the spots. The left figure highlights the total number of counts read for every gene, and the right figure summarises the number of genes with at least 1 count in a cell, highlighting the diversity of gene expression patterns spatially across the tissue.

### A.1 Description of Cell2Location deconvolution pipeline

The desired cell type compositions of the spots in Cell2Location are obtained using two main steps:

1. Computing cell type specific gene expression *signatures* using reference single-cell RNA-seq data.
2. Using variational inference to sample latent posterior distributions for the cell type proportions.

#### A.1.1 Computing reference cell type signatures

Cell type signatures are obtained by performing regularised Negative Binomial regression. The motivation behind using this model is that it would robustly derive the reference expression values of cell types *g*_*f,g*_ using input data composed of different batches *e* = {1, …, *E*} and technologies *t* = {1, …, *T*} that may affect the results (though for our case studies this is actually not utilised) (Kleshchevnikov et al., 2022). Here the expression count matrix *J* = {*j*_*c,g*_}, *c* ∈*C, g*∈ *G* follows a Negative Binomial distribution with unobserved expression levels (rates) *μ*_*c,g*_ and a gene-specific over-dispersion *α*_*g*_:

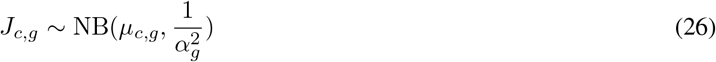

*μ*_*c,g*_ is modelled as a linear function of the reference cell type signatures and the batch/technical effects:

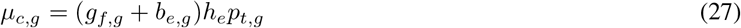

where *h*_*e*_ is a global scaling parameter between experiments *e* (for example difference in sequencing depth, or the number of times a given nucleotide has been read in an experiment (Lopez et al., 2018)). *p*_*t,g*_ accounts for multiplicative gene-specific difference in sensitivity between technologies, *b*_*e,g*_ accounts for additive background shift of each gene in each experiment *e* caused by free-floating RNA.

The priors of these variables are specified in hierarchical manner using similar constructions as we have seen previously with Cell2Location:

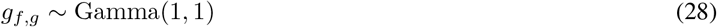

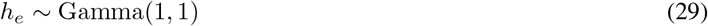

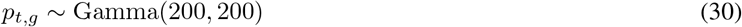

The prior on the additive shift variable *b*_*e,g*_ is specified in the same manner as *s*_*e,g*_ (Equations 7.21-7.25). The prior on *α*_*g*_ is specified similarly to *α*_*e,g*_ (Equations 7.18-7.20):

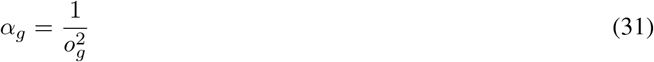

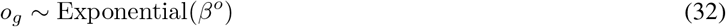

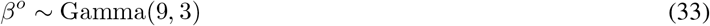

All model parameters are constrained to be positive. A weak L2 regularisation of *g*_*f,g*_, *b*_*e,g*_, and *α*_*g*_ is set alonside a strong penalty for deviations of *h*_*e*_ and *p*_*t,g*_ from 1. The average for each cell type *f* is used to initialise *g*_*f,g*_. *b*_*e,g*_ is initialised at the average expression of each gene *g* in each experiment *e* divided by 10 (Lopez et al., 2018; Kleshchevnikov et al., 2022).

#### A.1.2 Inference

Stochastic variational Bayesian inference is used to approximate the posterior distributions, enacted through Pyro (Bingham et al., 2018) and its autodiff variational inference framework (ADVI). Briefly, the latent posterior distributions of the model are approximated using univariate normal distributions which are softplus transformed to ensure positive scaling. The parameters of the variational distributions are chosen through minimisation of the KL divergence between the variational approximation and the true posterior distribution. This is equivalently achieved by maximising the evidence lower bound (ELBO objective).

After this optimisation, the posterior mean, standard deviation, 5% and 95% quantiles for each parameter are computed using 1000 samples from the variational posterior distribution. The mean was used for all of the results we show in the results section.

### A.2 GNN-C2L: spatially aware spatial cell deconvolution

Our methods build upon the Cell2Location pipelines, introducing relational inductive biases enacted by different instances of the message passing neural network framework. This primarily consists of two major additions. The first is the construction of the underlying graph that establishes proximal relationships between each of the observations we are interested in: the spots. The second is the augmentation of the Cell2Location generative model and the introduction of graph neural networks into them.

#### A.2.1 Constructing a spatial proximity graph on the spatial rna-seq output

As with many GRL methods, the underlying graph is an important factor behind actualising the assumption we intend to incorporate with relational inductive biases and obtaining performance gains (Hamilton). However, first and foremost we explore the utilisation of proximity and the assumption that proximal spots exhibit similar cell type compositions or specific cell type relationships between spots. To utilise this we want a graph neural network to operate on the proximity graph of spots after they have been selected by standard preprocessing pipelines (Heumos et al., 2023; Lopez et al., 2018; Kleshchevnikov et al., 2022). Depending on the positional arrangement of spots dictated by the spatial transcriptomics technology used — for example, hexagonal arrangement in 10x Visium slides as seen in Figure 4, or the grid arrangement found in our pseudo-synthetic dataset — a different number of neighbours is specified alongside the size of the local perceptive field we want a single layer of a GNN to operate over.

**Figure 4:**
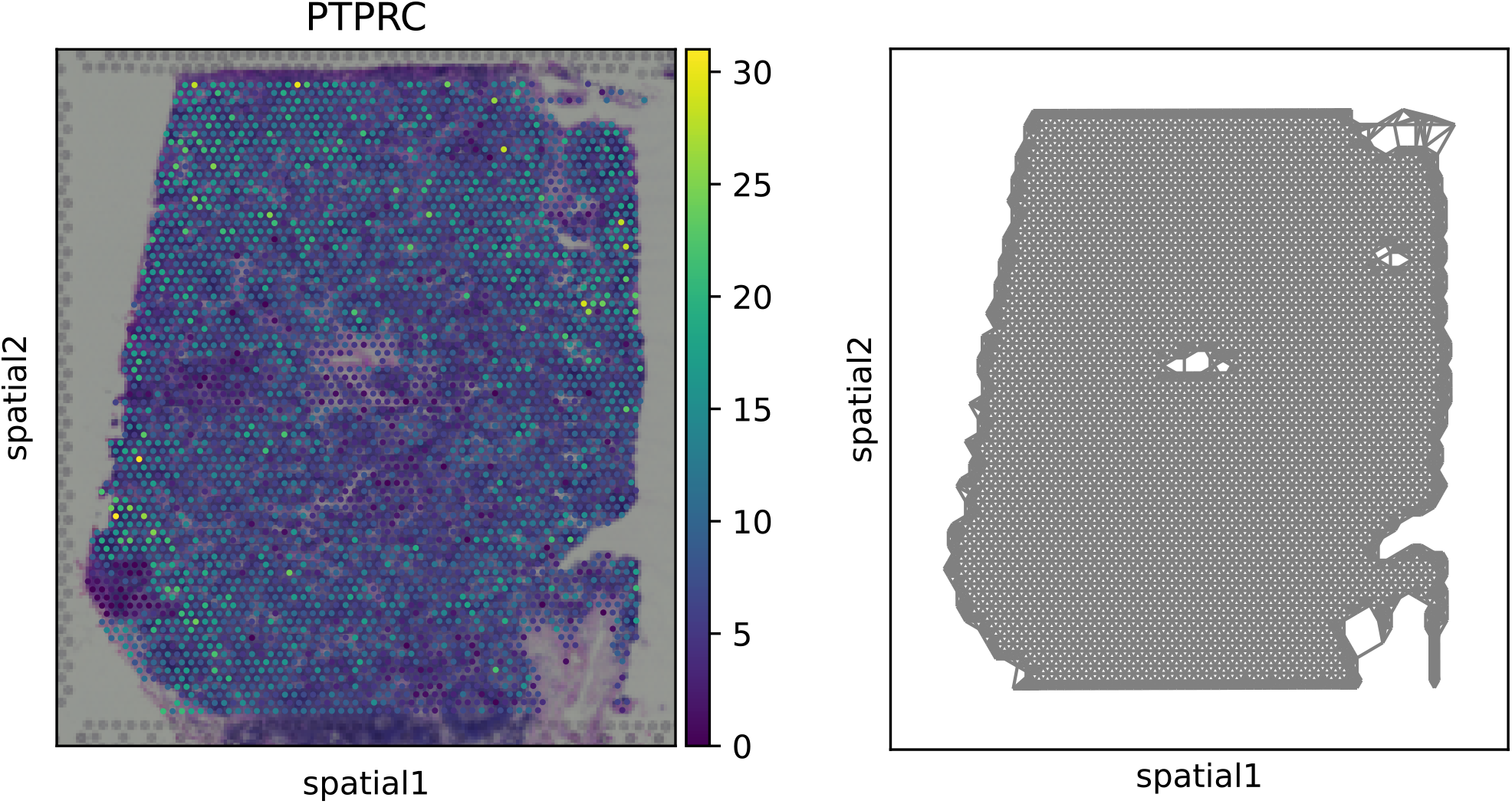
An example of the proximity graph computed using 6 neighbours for each spot in the hexagonal arrangement utilised in the 10X Visium protocol for a human lymph node sample (publicly available through the 10x Genomics dataset portal, accessed through scanpy (Wolf et al., 2018)). On the left is the output of the spatial transcriptomics sequencing with colouring of the spots based on the transcript counts for the gene PTPRC. The right shows the underlying proximity graph between each of the spots with its 6 closest neighbours.

#### A.2.2 GNN-C2L

We propose a hierarchical model for cell-type composition inference that incorporates proximal relationships between spots. Let *𝒩* (*s*) be the set of neighbour indices for spot *s*. This set of neighbours can be adapted to various spatial arrangements (e.g. hexagonal neighbourhoods for 10X Visium data) and k-hop neighbourhoods. To account for the neighbourhood information, we introduce a latent variable *γ*_*s,f*_ representing the neighbour-aware cell-type abundances:

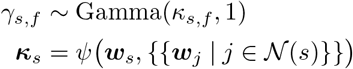

where the shape parameter *κ*_*s,f*_ depends on the latent variables ***w***_*s*_ and {{***w***_*j*_| *j ∈ 𝒩*(*s*)}} of spot *s* and its neighbours through a transformation *ψ*(·). Unlike Cell2Location, this effectively adds graphical dependencies between the neighbour-informed variables *γ*_*s,f*_ and the latent variables *w*_*s,f*_ (Appendix A). Importantly, computing *γ*_*s,f*_ as a function of ***w***_*s*_ allows capturing cell-type co-location patterns.

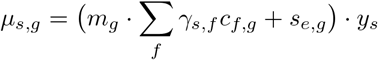

##### Incorporating spatial inductive biases

The form of *ψ*(·) determines the inductive biases of the model. In this study, we construct a proximity graph of spatially localised spots, i.e. we consider physically adjacent spots, allowing for different ·spatial arrangements (e.g. hexagonal neighbourhoods for 10X Visium data) and k-hop neighbourhoods. We consider several graph neural network architectures for *ψ*(·), starting with simple graph convolutional network (Wu et al., 2019) to validate whether homophily (enacted by feature propagation) is a useful inductive bias, and introducing other GNN operators to allow for a more expressive use of the available spatio-relational data. We also consider a standard multi-layer perceptron as a baseline to assess whether performance changes can be attributed to similarly parametrised spatial-agnostic transformations. We next describe the alternatives for *ψ* in greater detail.

##### MLP-C2L

As a spatial-agnostic control, we model *ψ*(·) with an MLP, i.e. ***κ***_*s*_ = MLP(***w***_*s*_), using a softplus activation function. This model does not utilise any spatial relationships between the spots and, alongside Cell2Location, serves as a control for our hypothesis. |N |

##### SGC-C2L

We construct a GNN-C2L variant using Simple Graph Convolutional (SGC) layers (Wu et al., 2019; Scherer et al., 2019). Let *d*_*s*_ = | *𝒩* (*s*)| be the node degree of spot *s*. A SGC layer computes the neighbour-aware features ***κ***_*s*_ using a weighted average of the latent variables ***w***_*s*_ in the local neighbourhood

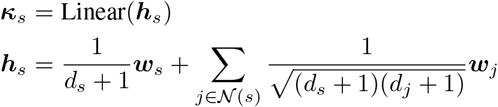

##### GAT-C2L

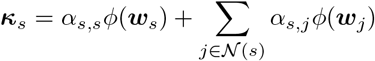

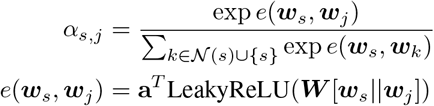

## B. Dataset details

### B.1 Purely synthetic dataset

Briefly, this simulated spatial transcriptomic dataset introduced in Cell2Location (Kleshchevnikov et al., 2022) consists of 2,500 spots by sampling cells from 49 reference cell types, using a combination of ubiquitous abundance patterns and cell types distributed according to regional tissue zones as observed in semi-simulated data (Kleshchevnikov et al., 2022). Reference cell type signatures were adapted from a mouse brain scRNA-seq dataset sequenced by Kleshchevnikov et al. (2022). The dataset reflects diverse cell abundance patterns across ubiquitous and spatially restricted cell types, which allow us to evaluate our methods within different cell abundance scenarios in addition to just overall accuracy. The paired reference scRNA-seq dataset consists of 8,111 cells exhibiting 12,080 genes whose expression is measured (recall this number is an intersection of the set of transcripts measured in the scRNA-seq and spatial transcriptomic experiments). This dataset is preprocessed and utilised to compute reference cell type signatures matrix *𝒞* as detailed in Appendix C.1. More information on the construction of this dataset is detailed extensively in Section 5 of the supplementary materials in (Kleshchevnikov et al., 2022).

### B.2 MPOA

Briefly, the MPOA dataset is a simulated spatial transcriptomic dataset constructed by aggregating the gene expression of cells from single-cell resolution multiplex error robust fluorescence in situ hybridisation (MERFISH) data of the mouse medial preoptic area (Moffitt et al., 2018). The availability of both single-cell resolution gene expression in-situ makes it an ideal candidate for creating simulated spatial transcriptomic data. It contains measurements of 135 genes selected to distinguish between major non-neuronal cell types as well as neuronal subtypes. Transcriptional clustering analysis on the scRNA-seq measurements identified 9 major cell types which are used to form the underlying target proportions we wish to infer. To simulate the multi-cellular spatial transcriptomic data, the single-cell resolution MERFISH data is aggregated into 100*μm*^2^ pixels which we proxy as spot transcripts. More information on this dataset and its availability can be found in the supplementary materials in (Miller et al., 2022).

### B.3 Xenium

The Xenium breast cancer dataset (Janesick et al., 2022) jointly profiled the expression and spatial location of 167,885 cells (313 genes) from a formalin-fixed, paraffin-embedded (FFPE) human breast cancer section. We utilised squidpy (Palla et al., 2022) to download, process the data, and detect Leiden communities (https://squidpy.readthedocs.io/en/stable/external_tutorials/tutorial_xenium.html). We then employed decoupler-py (Badia-i Mompel et al., 2022) to perform overrepresentation analysis (Figure 5), using breast cancer marker genes (Janesick et al., 2022). In total, we identified 8 broad cell types including immune cells (e.g. T cells, B cells, and macrophages) and breast cells (e.g. grandular cells, myoepithelial cells, and cancer cells). We aggregated (i.e. summed) the single-cell resolution Xenium data into 100*μm*^2^ pixels as a proxy for convolved spot transcripts.

**Figure 5:**
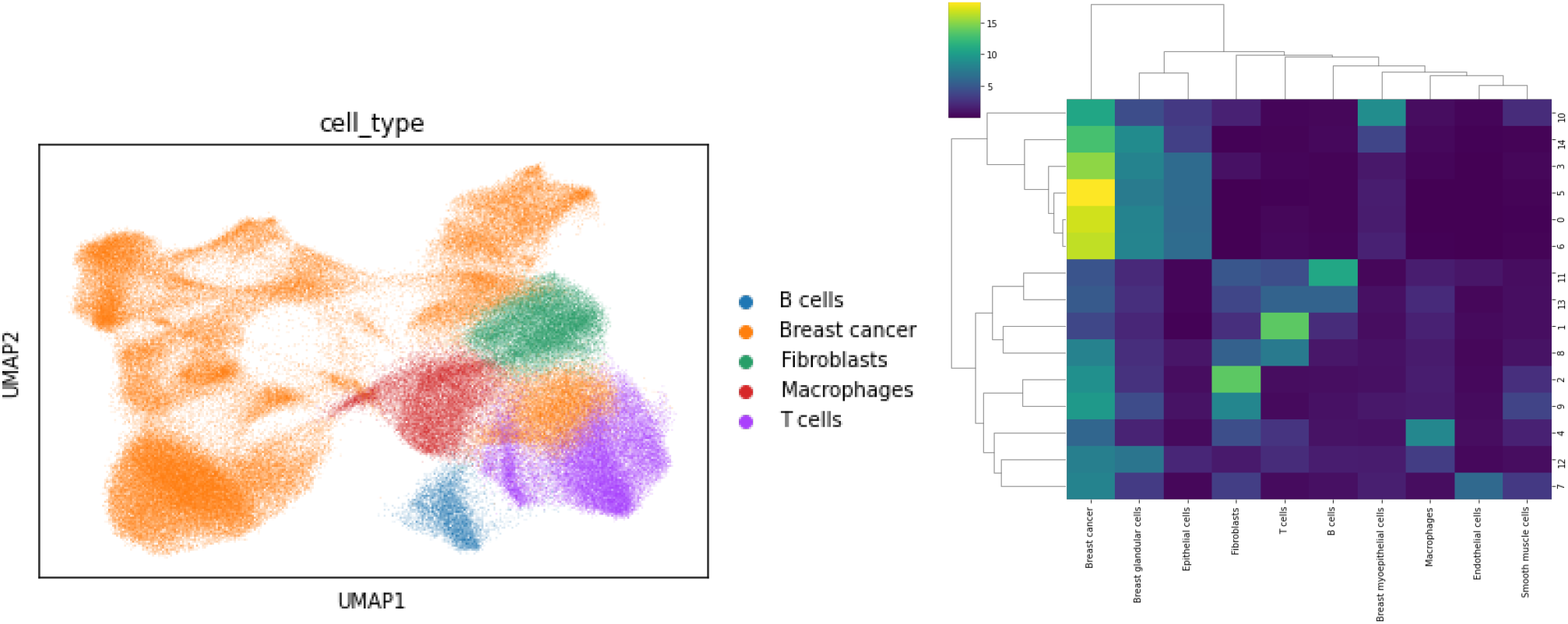
(left) UMAP plot of Xenium colored by Leiden clusters. (right) Overrepresentation analysis scores for each Leiden cluster.

## C. Additional experimental setup details

### C.1 Inference of cell type signatures

As in Cell2Location (Kleshchevnikov et al., 2022), two approaches can be taken to compute cell type signatures matrix ***C*** ∈ ℝ^*F* ×*G*^:

#### 1. Inference using Negative Binomial regression

Given the raw untransformed and unnormalised count matrix **J∈**ℝ^|*C*|×|*G*|^ of the reference scRNA-seq data, several preprocessing steps were taken before it is input to the NB regression to find cell type signatures. The data is filtered at 2 cut-offs: i) selecting genes detected that have more than 0 mRNA counts for at least 5% of the cells, and ii) selecting genes with a mean expression greater than 1.1 and a greater than 0 mRNA count in at least 0.05% of the cells. Subsequently it is given to the NB regression algorithm to infer the cell type signatures.

To infer the cell type signatures, the variational parameters of the Negative Binomial regression model were trained with stochastic gradient descent on the ELBO objective using a batch size of 1024 cells and an Adam optimiser with a learning rate set at 0.001 for 500 epochs. The posterior cell signature values were sampled 1000 times and the mean values were used. To control and simplify comparison between the deconvolution models, the cell type signatures are shared between each of the methods we evaluate.

This approach is used in the purely synthetic dataset, and is typically the recommended approach in Cell2Location as the regression model is robust to different technology and batch effects.

#### 2. Using mean counts of genes for each cell type

Given the raw untransformed and unnormalised count matrix **J∈**ℝ^|*C*|×|*G*|^ of the reference scRNA-seq data, we compute the average count of genes *g ∈ G* for each of the cell types in *F* to construct ***C*∈*ℝ***^*F* ×*G*^. This approach is computationally efficient and can provide good mappings when the scRNA-seq data is obtained from a single batch or better yet is paired with the spatial transcriptomic data, as is the case in the MPOA and Xenium datasets. This approach was utilised in our evaluation for MPOA and Xenium.

## D. Effect of increasing neighbourhood size

The increased performance through the utilisation of the 1-hop proximal neighbourhoods begs the question of how the size of the receptive field can influence performance. Recall from Section 2.4.2 that this is as simple as adding more layers to the graph neural network layers. We present the table of results examining increase of receptive field with SGC-C2L and GAT-C2L in Tables 2, 3 and 4. Whilst all of the models still consistently outperform the original Cell2Location and MLP-C2L, we can see certain performs patterns that are in line with GNN theory. Specifically, in all but the average JSD metric for ULCA we see that the best performing models exist in the first models exhibiting up to 3 layers, exhibiting a drop in performance after the best performance. This pattern is common GNN based methods due to the oversmoothing and oversquashing phenomenon (Alon & Yahav, 2020) that prevents GNNs from effectively incorporating information from distant neighbours as the aggregation of messages into fixed size vectors creates an information bottleneck. In addition to the mechanical limitation of the GNNs, we also have to consider the relationship between the growing receptive field and its absolute distance away from the target spot we want to influence in terms of the sizes of cell colocation patterns. Depending on the cell-types, tissue architecture, and fidelity of the spatial transcriptomic technology, differing receptive field sizes over spots will be biologically relevant to capturing cell colocation patterns.

**Table 2:**
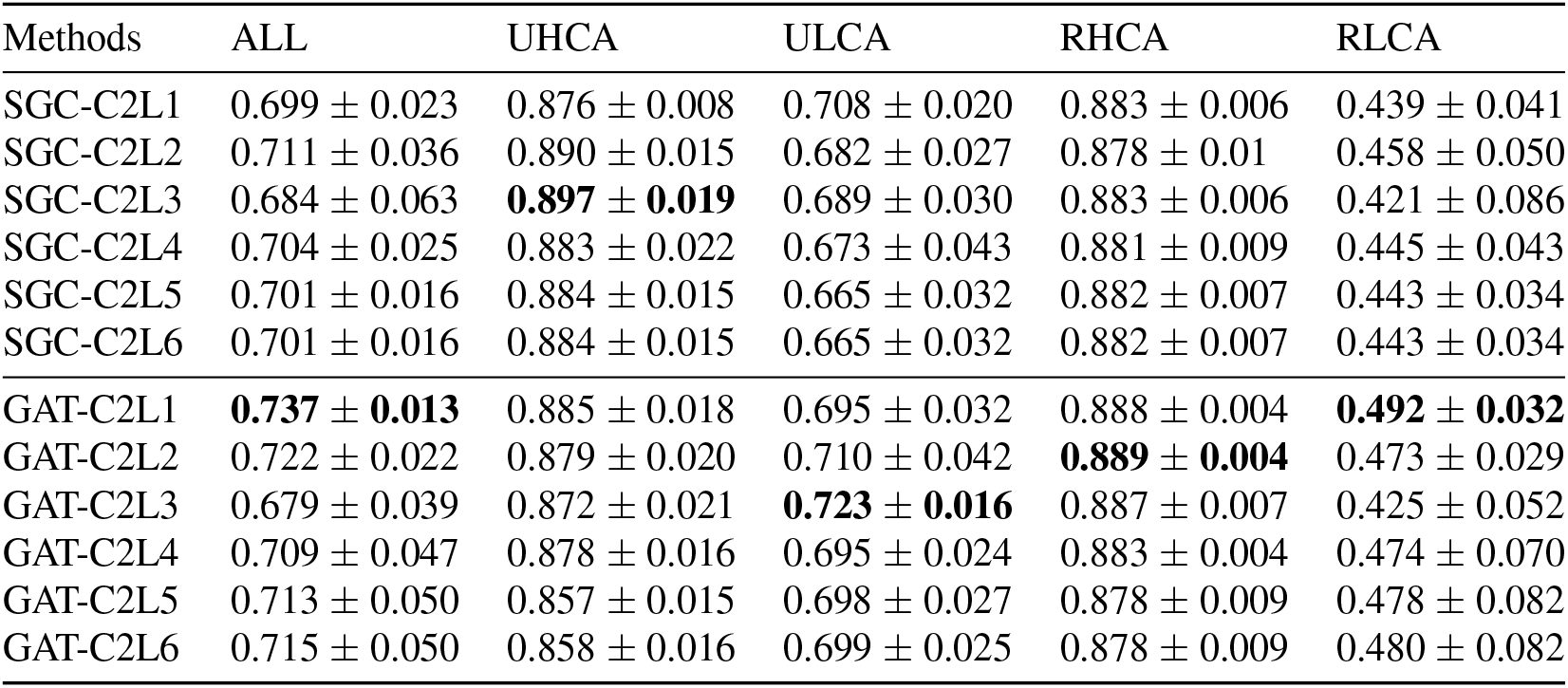
Average Pearson R correlation and standard deviation of 5 seeded runs of each model over all spots. Correlation values for subcategories of cell types exhibiting distinct cell abundance patterns are also provided. Bold numbers indicate best performing method for each category of cell types being evaluated.

| Methods | ALL | UHCA | ULCA | RHCA | RLCA |
| --- | --- | --- | --- | --- | --- |
| SGC-C2L1 | 0.699 ± 0.023 | 0.876 ± 0.008 | 0.708 ± 0.020 | 0.883 ± 0.006 | 0.439 ± 0.041 |
| SGC-C2L2 | 0.711 ± 0.036 | 0.890 ± 0.015 | 0.682 ± 0.027 | 0.878 ± 0.01 | 0.458 ± 0.050 |
| SGC-C2L3 | 0.684 ± 0.063 | **0.897 ± 0.019** | 0.689 ± 0.030 | 0.883 ± 0.006 | 0.421 ± 0.086 |
| SGC-C2L4 | 0.704 ± 0.025 | 0.883 ± 0.022 | 0.673 ± 0.043 | 0.881 ± 0.009 | 0.445 ± 0.043 |
| SGC-C2L5 | 0.701 ± 0.016 | 0.884 ± 0.015 | 0.665 ± 0.032 | 0.882 ± 0.007 | 0.443 ± 0.034 |
| SGC-C2L6 | 0.701 ± 0.016 | 0.884 ± 0.015 | 0.665 ± 0.032 | 0.882 ± 0.007 | 0.443 ± 0.034 |
| GAT-C2L1 | **0.737 ± 0.013** | 0.885 ± 0.018 | 0.695 ± 0.032 | 0.888 ± 0.004 | **0.492 ± 0.032** |
| GAT-C2L2 | 0.722 ± 0.022 | 0.879 ± 0.020 | 0.710 ± 0.042 | **0.889 ± 0.004** | 0.473 ± 0.029 |
| GAT-C2L3 | 0.679 ± 0.039 | 0.872 ± 0.021 | **0.723 ± 0.016** | 0.887 ± 0.007 | 0.425 ± 0.052 |
| GAT-C2L4 | 0.709 ± 0.047 | 0.878 ± 0.016 | 0.695 ± 0.024 | 0.883 ± 0.004 | 0.474 ± 0.070 |
| GAT-C2L5 | 0.713 ± 0.050 | 0.857 ± 0.015 | 0.698 ± 0.027 | 0.878 ± 0.009 | 0.478 ± 0.082 |
| GAT-C2L6 | 0.715 ± 0.050 | 0.858 ± 0.016 | 0.699 ± 0.025 | 0.878 ± 0.009 | 0.480 ± 0.082 |

**Table 3:** Average of average Jensen-Shannon divergence (JSD) along with standard deviation of 5 seeded runs of each model. JSD values for subcategories of cell types exhibiting distinct cell abundance patterns are also provided. Bold numbers indicate best performing method for each category of cell types being evaluated.

| Methods | ALL | UHCA | ULCA | RHCA | RLCA |
| --- | --- | --- | --- | --- | --- |
| SGC-C2L1 | $0.446 \pm 0.006$ | $0.224 \pm 0.011$ | $0.460 \pm 0.007$ | $0.368 \pm 0.005$ | $0.493 \pm 0.009$ |
| SGC-C2L2 | $0.443 \pm 0.007$ | $0.208 \pm 0.021$ | $0.467 \pm 0.010$ | $0.371 \pm 0.009$ | $0.489 \pm 0.007$ |
| SGC-C2L3 | $0.447 \pm 0.011$ | **$0.199 \pm 0.017$** | $0.463 \pm 0.006$ | $0.369 \pm 0.007$ | $0.499 \pm 0.015$ |
| SGC-C2L4 | $0.448 \pm 0.006$ | $0.216 \pm 0.019$ | $0.472 \pm 0.014$ | $0.375 \pm 0.008$ | $0.494 \pm 0.009$ |
| SGC-C2L5 | $0.448 \pm 0.005$ | $0.207 \pm 0.022$ | $0.473 \pm 0.010$ | $0.375 \pm 0.007$ | $0.493 \pm 0.008$ |
| SGC-C2L6 | $0.448 \pm 0.005$ | $0.207 \pm 0.022$ | $0.473 \pm 0.010$ | $0.375 \pm 0.007$ | $0.493 \pm 0.008$ |
| GAT-C2L1 | **$0.435 \pm 0.003$** | $0.209 \pm 0.021$ | $0.458 \pm 0.014$ | $0.369 \pm 0.001$ | **$0.482 \pm 0.006$** |
| GAT-C2L2 | $0.438 \pm 0.006$ | $0.223 \pm 0.017$ | $0.458 \pm 0.014$ | $0.363 \pm 0.002$ | $0.486 \pm 0.005$ |
| GAT-C2L3 | $0.447 \pm 0.008$ | $0.222 \pm 0.025$ | $0.450 \pm 0.009$ | **$0.356 \pm 0.004$** | $0.496 \pm 0.011$ |
| GAT-C2L4 | $0.441 \pm 0.010$ | $0.215 \pm 0.018$ | $0.452 \pm 0.010$ | $0.358 \pm 0.004$ | $0.487 \pm 0.013$ |
| GAT-C2L5 | $0.445 \pm 0.015$ | $0.243 \pm 0.017$ | **$0.448 \pm 0.009$** | $0.362 \pm 0.007$ | $0.492 \pm 0.020$ |
| GAT-C2L6 | $0.444 \pm 0.015$ | $0.242 \pm 0.017$ | $0.448 \pm 0.009$ | $0.362 \pm 0.007$ | $0.491 \pm 0.020$ |

**Table 4:** Average AUPRC scores and standard deviation of 5 seeded runs of each model over all spots. Scores for subcategories of cell types exhibiting distinct cell abundance patterns are also provided. Bold numbers indicate best performing method for each category of cell types being evaluated.

| Methods | ALL | UHCA | ULCA | RHCA | RLCA |
| --- | --- | --- | --- | --- | --- |
| SGC-C2L1 | $0.719 \pm 0.002$ | $0.977 \pm 0.004$ | $0.646 \pm 0.006$ | $0.861 \pm 0.001$ | $0.719 \pm 0.002$ |
| SGC-C2L2 | $0.716 \pm 0.003$ | $0.978 \pm 0.001$ | $0.644 \pm 0.006$ | $0.860 \pm 0.001$ | $0.716 \pm 0.003$ |
| SGC-C2L3 | $0.710 \pm 0.002$ | **$0.979 \pm 0.002$** | $0.649 \pm 0.005$ | $0.852 \pm 0.001$ | $0.710 \pm 0.002$ |
| SGC-C2L4 | $0.701 \pm 0.004$ | $0.972 \pm 0.003$ | $0.639 \pm 0.007$ | $0.845 \pm 0.005$ | $0.701 \pm 0.004$ |
| SGC-C2L5 | $0.701 \pm 0.007$ | $0.975 \pm 0.003$ | $0.633 \pm 0.009$ | $0.848 \pm 0.005$ | $0.701 \pm 0.007$ |
| SGC-C2L6 | $0.701 \pm 0.007$ | $0.975 \pm 0.003$ | $0.633 \pm 0.009$ | $0.848 \pm 0.005$ | $0.701 \pm 0.007$ |
| GAT-C2L1 | $0.722 \pm 0.002$ | $0.978 \pm 0.004$ | $0.664 \pm 0.004$ | $0.858 \pm 0.003$ | $0.722 \pm 0.002$ |
| GAT-C2L2 | **$0.726 \pm 0.001$** | $0.977 \pm 0.003$ | $0.665 \pm 0.007$ | $0.865 \pm 0.001$ | **$0.726 \pm 0.001$** |
| GAT-C2L3 | $0.721 \pm 0.003$ | $0.970 \pm 0.003$ | **$0.679 \pm 0.006$** | **$0.870 \pm 0.002$** | $0.721 \pm 0.003$ |
| GAT-C2L4 | $0.710 \pm 0.003$ | $0.968 \pm 0.002$ | $0.670 \pm 0.006$ | $0.867 \pm 0.001$ | $0.710 \pm 0.003$ |
| GAT-C2L5 | $0.700 \pm 0.002$ | $0.959 \pm 0.001$ | $0.652 \pm 0.010$ | $0.865 \pm 0.001$ | $0.700 \pm 0.002$ |
| GAT-C2L6 | $0.702 \pm 0.003$ | $0.961 \pm 0.003$ | $0.652 \pm 0.009$ | $0.865 \pm 0.001$ | $0.702 \pm 0.003$ |

